# Size-Based Dominance Hierarchy In One Of Two Sympatric Cryptic Pacific Skinks (*Emoia impar* And *Emoia Cyanura*)

**DOI:** 10.1101/2020.01.11.902866

**Authors:** Mary “Molly” Hallsten

## Abstract

*Emoia impar* and *Emoia cyanura* are two morphologically cryptic Pacific skinks that have different preferred thermal micro-habitats but similar geographic range and overlap. Previously individuals have been noted to display a size-based dominance hierarchy at favored basking sites, though this behavior was not specified between species. I found that only one of the two species, *E. impar*, naturally presents this size-based dominance hierarchy in areas of high population density. Neither species exhibit the hierarchy in low population density areas. No evidence was found to suggest that the presence of this hierarchy allows one species to exclude the other.

## INTRODUCTION

Understanding how multiple species co-exist within a shared habitat is a fundamental challenge in ecology (Gause 1934). Hardin (1960) documented the competitive exclusion principle, which states that if two non-interbreeding populations occupy the same ecological niche and geographical territory, and differ in their rate of resource consumption, one species will eventually drive the other to extinction. Therefore, for two species to co-exist, there must be some difference in either behavior or exploitation of resources. An aspect of co-existence is resource partitioning, where two species differ in the use of a resource within the same fundamental niche (Vandermeer 1974). In ectotherms, one aspect of niche partitioning may come in the form of thermal regulation and physiological differences (Gunderson et al. 2018).

Thermal regulation and the availability of favored basking sites can create competition between individuals (Lewis et al. 2000). For example, Lewis (2000) notes the presence of size-based dominance hierarchies in a species of whiptail, *Pholidoscelis exsul*, where the males exhibit aggressive interactions when they compete for basking and foraging sites in an experimental setting. Dominance hierarchies may be expected to occur in habitats of high population density due to an increased frequency of aggressive interactions (Alberts 1994; Brattstrom 1974; Toft 1985). Brattstrom (1974) claimed that reptilian hierarchies are often a result of experimentally crowded territorial species, where an individual’s size determines their position in the hierarchy. Less is known about naturally occurring dominance hierarchies, particularly amongst members of the family Scincidae.

*Emoia impar* and *Emoia cyanura* are two morphologically cryptic skink species located throughout much of Oceania, from the Solomon Islands to French Polynesia and Micronesia with broad range overlap (Zug 2013). Bruna et al. (1996) determined the two species to be morphologically convergent but not sister taxa, and that they co-existed in all habitats studied. They partition micro-habitats by thermal preference, with *E. cyanura* preferring open canopy cover, a higher body temperature, and warmer substrates than *E. impar* (McElroy 2014).

The behavioral differences that allow for *E. impar* and *E. cyanura* to coexist is not yet fully understood. A diet study on the island of Mo’orea by Erica Garcia (1997) observed *E. impar* consuming four prey categories that *E. cyanura* did not, but the study did not indicate that foraging or diet significantly impacts the distribution of either species. Other than diet, a thermal or behavioral distinction should be considered. Zug (1991) recorded that *E. impar* and *E. cyanura* are behaviorally similar. Zug also noted that when frightened away, skinks of both species would recolonize sunlit areas with the smallest individuals appearing first, followed by the successive displacement of smaller individuals by increasingly larger ones. These observations suggest the presence of a dominance hierarchy, but Zug did not differentiate between species to note if this was intraspecific competition or interspecific exclusion from optimal sunning sites.

Both *E. impar* and *E. cyanura* are frequently and easily observed on the island of Mo’orea, French Polynesia, suggesting that both are abundant. Given the high number of skinks seen with preliminary observations, the abundance of these two *Emoia* skinks on Mo’orea is part of what makes the island a model system to study their interactions. On islands of Fiji, the micro-habitat of *E. cyanura* tends to overlap with other species beyond *E. impar*, such as *Gehyra oceanica, Hemiphyllodactylus typus, Lepidodactylus lugubris, Nactus pelagicus, Cryptoblepharus eximius, Emoia concolor, Emoia nigra, Emoia trossula, Lipinia noctua*, and *Leiolopisma alazon* (Zug 1991). In contrast, the island of Mo’orea is home to only four species of skinks: *Lipinia noctua, Cryptoblepharus poecilopleurus, Emoia impar*, and *Emoia cyanura*, with the *Emoia* skinks being the two most observable and the two with the closest observed overlapping distributions. Mo’orea is an ideal island to observe interactions between these two Emoia species with minimal interference from other herpetofauna.

A study of *E. impar* and *E. cyanura* was undertaken to investigate behavioral interactions between individuals at basking sites in varying habitats on the island of Mo’orea, and how these interactions relate to their co-existence. This study aims to examine possible hierarchies present between individuals of the same species as well as between species, while also investigating if basking hierarchies result in one species excluding the other from shared habitat. I hypothesize that this size-based dominance hierarchy is present in both species and allows for individuals to exclude members of the opposite species. I also hypothesize that this behavior will be present only in densely populated sites where competition is greatest.

## MATERIALS AND METHODS

### Study sites

This study was conducted on the island of Mo’orea, French Polynesia throughout the Opunohu Valley, including and along the Three Coconuts Trail (Fig. 1). Sites were selected if four or more skinks were initially observed regardless of species.

**FIG. 1.**
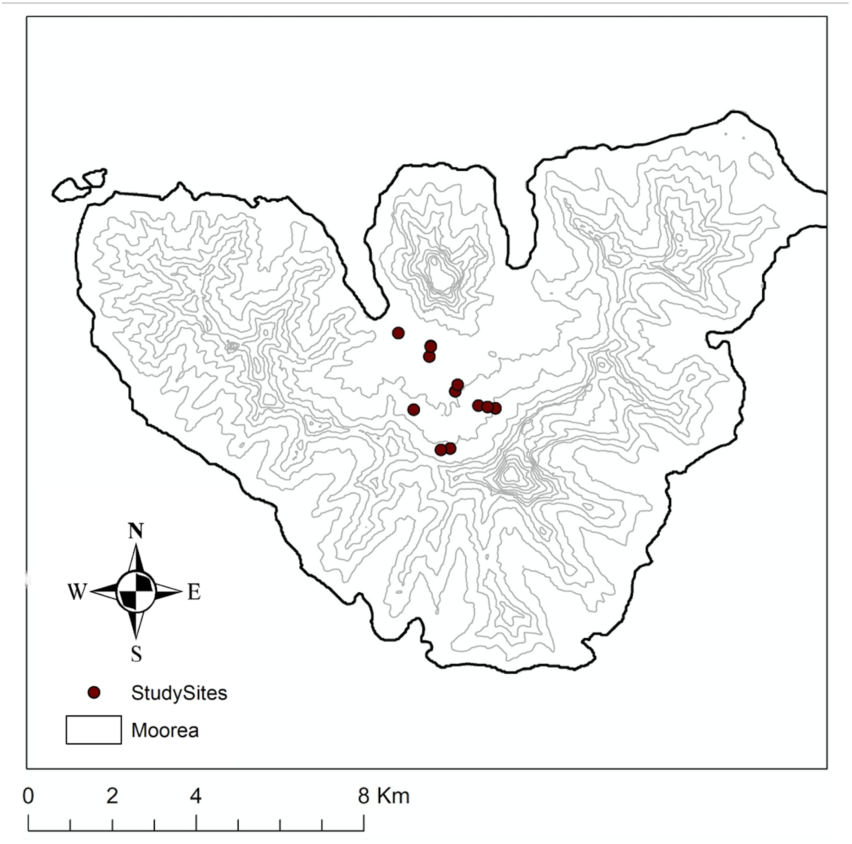
Map, Mo’orea, French Polynesia showing location of study sites (Table 1A in Appendix)

### Field experiment

To explore this naturally occurring basking hierarchy, I conducted a field experiment. After locating a site with at least four skinks, I chased the skinks away and started a 10-minute timer. Photographs, time of re-appearance, order of re-appearance, and basking substrate were recorded for each individual that recolonized the site within the 10-minute period. Order of re-appearance is analogous with order in hierarchy for all graphs and analysis. After the 10-minutes, basking substrate dimensions were taken to serve as scales in the photographs. I imported the photos to the program ImageJ (Schneider et al. 2012), where I measured the length of each skink from snout to vent relative to the scale bar in each photograph. All photographs were taken with a Nikon Coolpix P500. A total number of skinks seen before, during, and after the observation period was taken.

### Identifying species

Species of each individual was determined based on the following three characteristics: (1) blue or green tail coloration, (2) absence or presence of parietal eye, and (3) if the middorsal scales fused into larger single scales (*E. impar*) or remained unfused (*E. cyanura*). (Ineich and Zug 1991). Photographs and binoculars were used to closely examine every individual to ensure correct identification. Photographs were also closely examined during data analysis to ensure one individual was not counted more than once per site. Identification guides are available in the Appendix (Figures 5A, 6A).

### Sites variation

After each observational period, ten thermal measurements of the varying sunning substrates were taken with an Infrared Thermometer [Fischer Scientific] and averaged. One quadrat varying between 2×2 or 3×3 meters was taken at each site and the following measurements were recorded in percentages of cover: sunlight, wood, shrubs, soil, rocks, and leaf litter. Percent of canopy cover was recorded at each site using the iPhone application “Percentage Cover” (Public Interest Enterprises 2017). Number of trees were recorded, with a tree being categorized as any woody plant over one meter in height within the quadrat. GPS coordinates and elevation were recorded with the iPhone application “Altimeter” (CMH Digital 2019). Sites were categorized as either “high” populations or “low” populations. High population sites were sites where 10 or more total individuals of either species were observed.

### Statistical analysis

The relationship between hierarchy order and size of individual was analyzed using linear regression. The correlation between order and size was examined separately at both low population sites and high population sites.

Differences in mean between the first and last individuals at all sites were analyzed with a paired t-test. Tests were performed separately for each species. *Emoia impar* was analyzed separately at high and low population sites, whereas *E. cyanura* could only be examined at low population sites due to a low number of *E. cyanura* individuals at high population sites.

Habitat variables were compared with total skinks observed at a site using linear regression. Habitat variables were compared to one another using linear regression.

All analyses were done in R Studio (R Studio Team 2016). Alpha = 0.05 for all analysis.

## RESULTS

### Field experiment

The presence of a size-based dominance hierarchy was observed at high population sites among *E. impar* individuals, but not *E. cyanura. Emoia cyanura* did not exhibit this hierarchy behavior at low population sites (Figure 2c). Only 3 *E. cyanura* were observed across all high population sites, which was not a large enough sample size to analyze. *E. impar* displayed a size-based dominance hierarchy among the first four individuals to recolonize a high population basking site (Figure 2b), with smaller individuals appearing first before being subsequently replaced by larger individuals (p=.02, R^2^=.28). Analysis was done using linear regression. This behavior was not exhibited by *E. impar* at low population sites (p=0.98) (Figure 2a).

**FIG. 2.**
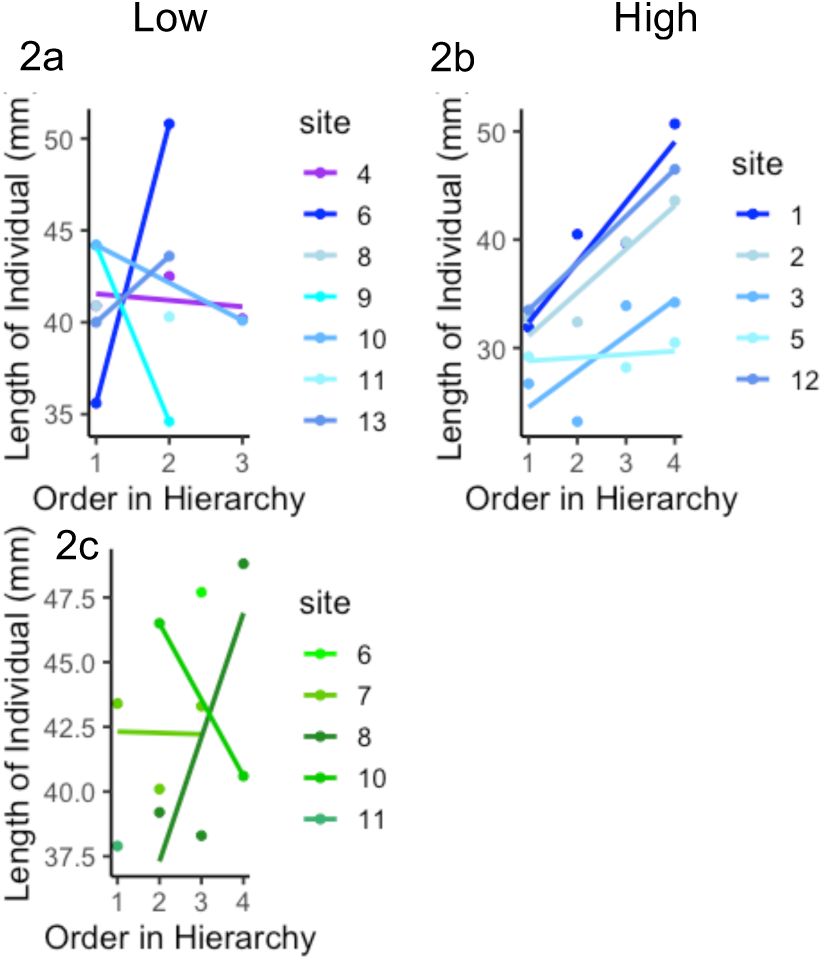
Relationship between body size and order in hierarchy. No observed hierarchy at low populations (left column) for either species (*E. cyanura*, green(2c); *E. impar*, blue(2a)). A significant correlation between order in hierarchy and length of individual (p<.02) for *E. impar* at high population sites (2b), with the smallest individuals recolonizing first, followed by larger individuals.

The mean length of the first and last individuals to recolonize the sun spot was compared between low population and high population sites with paired two-way t-tests. Difference of medians are shown for visualization (Figure 3). Species were analyzed separately. At high population sites, there was a significant difference in means between the first and last individuals of *E. impar* to recolonize a site (p<.05), (Figure 3b). Not enough *E. cyanura* individuals were observed at high population sites to analyze. At low population sites, there was no significant difference in means between the first and last individual for either species (*E. cyanura*, p = 0.63; *E. impar*, p = 0.84).

**FIG. 3.**
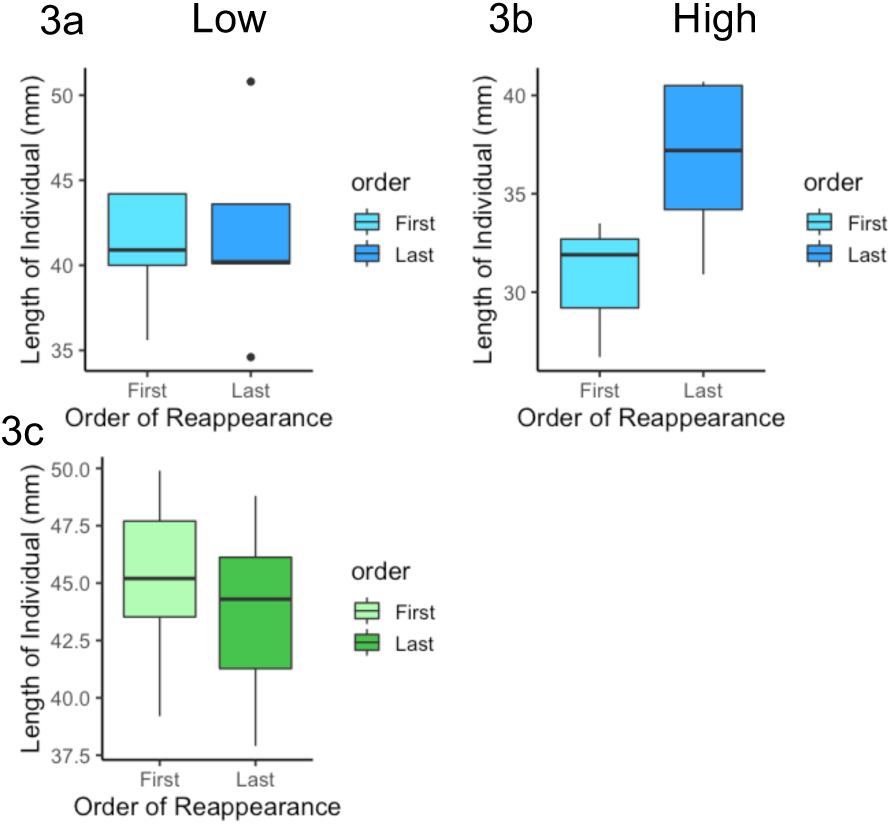
Relationship between body size and order of reappearance. Medians visualized in plot, means analyzed for p-values. No significant difference in mean length of first or last individual of either species at low population (*E. impar* 3a, *E. cyanura* 3c). Significant difference in means between first and last individual of *E. impar* (blue) at high population sites (3b) (p<.05). No high population data for *E. cyanura* (green) available for analysis.

### Sites variation

A total of 53 individuals were observed across 13 sites. Of these, 17 were *E. cyanura* and 36 of them were *E. impar*. Using linear regression, the number of individuals observed at a site significantly correlated at the p<.05 level with the following habitat characteristics: percent wood, percent leaf litter, percent shrubs, and substrate temperature. As noted in Table 2, higher populations correlated with lower percent wood cover (p<.001), lower percent shrub cover (p=.024), and relatively lower substrate temperatures (p=.001). High populations correlated positively with high percent leaf litter (p<.001). The only two habitat variables that correlated significantly (p<.05) were wood and leaf litter. Percent wood and percent leaf litter had a significant inverse relationship (p<.05).

**TABLE 2.** Correlation of habitat variables with total number of skinks observed at a sight. High percent leaf litter correlated positively with a high number of individuals observed. High percent wood cover, shrubs, and substrate temperature correlated negatively with number of individuals observed.

| Habitat Variable Percent Cover | Correlation | p-value | R<sup>2</sup> |
| --- | --- | --- | --- |
| Wood | Negative | <.001 | .21 |
| Leaf Litter | Positive | <.001 | .36 |
| Shrubs | Negative | .024 | .08 |
| Rock | Negative | .055 | .05 |
| Substrate Temperature | Negative | .001 | .17 |
| Sunspot Size | None | .606 | -.01 |

Substrate preference is displayed in Figure 4. Leaf litter was the most common basking substrate across both species. Only *E. impar* utilized live leaves and trees, whereas only *E. cyanura* utilized bamboo, though the sample size for these substrates was small.

**FIG. 4.**
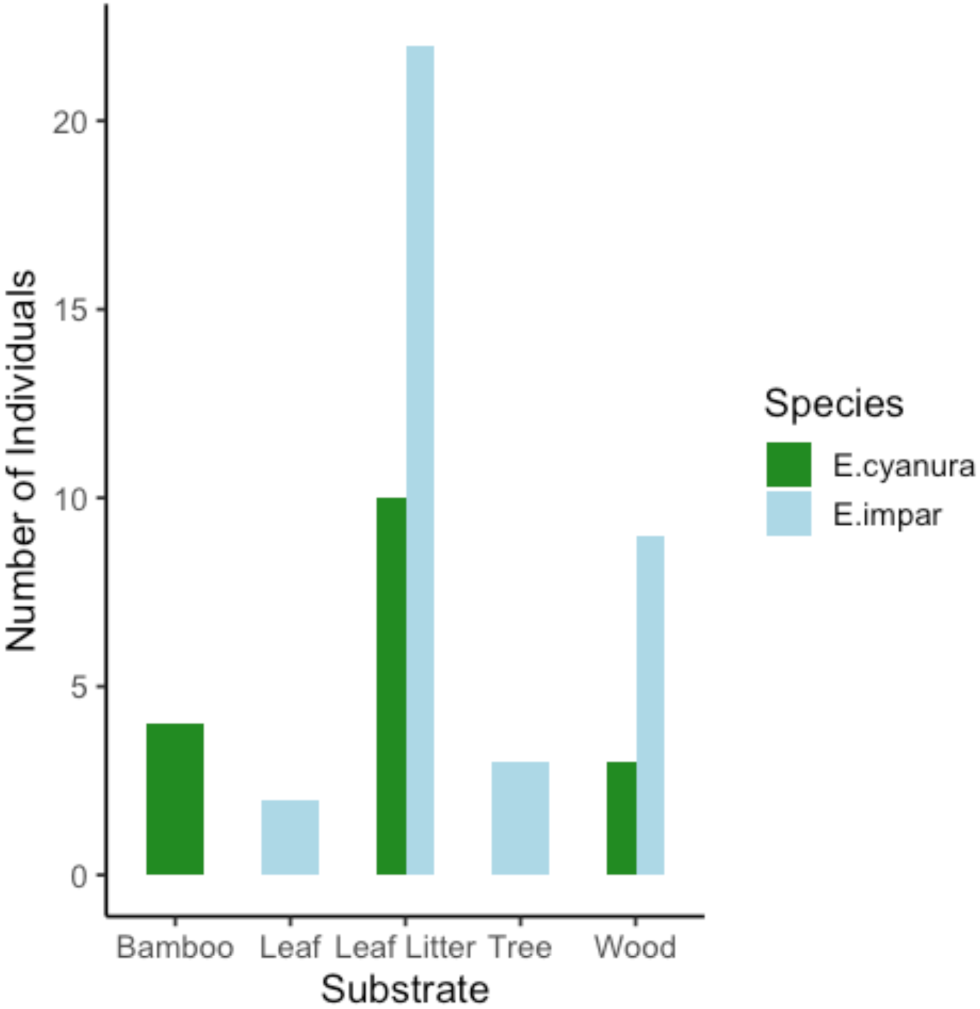
Abundance of individuals by substrate type. The most common basking substrate was leaf litter for both species. Only *E. cyanura* utilized bamboo, and only *E. impar* utilized live trees and leaves.

## DISCUSSION

My results show that an abundance of basking individuals correlates positively with a high percent ground cover of leaf litter (Table 2). Similarly, the higher percentage of wood ground cover or shrubs, the lower the number of observed basking individuals. These correlations may be attributed to both unintentional bias in methods as well as skink behavior. The presence of substrates that also serve as hiding spots such as wood logs or shrubs may provide skinks an area to remain hidden after being frightened off, therefore making them less likely to return to the observation site and resulting in a lower number of observed individuals. Individuals hiding amongst shrubs or deep in a wood pile may also exploit gaps in their canopy cover that allow for them to remain hidden while reaching their preferred temperature. These individuals would then be less likely to risk exposing themselves and I would not know to record them. Also, there are less hiding spots in sites that consist primarily of leaf litter, therefore individuals are more likely to move into my field of view when moving to the favored basking site. The significant inverse relationship between wood and leaf litter is a bias of my methods – it was impossible for a site to have both high percent wood and high percent leaf litter in my quadrats, for wood covering leaf litter was considered percent wood.

Substrate temperature correlated negatively with total individuals seen. As substrates increased in temperature later in the day, individuals were less likely to be observed sunning or remaining in one place. It is possible for lizards to overheat, so individuals were more likely to be foraging or hiding from direct sunlight when the temperature was too hot.

My hypothesis that a size-based dominance hierarchy is present among individuals in highly populated sites is correct for *E. impar*, but the same claim cannot be made about *E. cyanura* (Figure 3c). A size-based dominance hierarchy is not present among individuals of either species in areas with low competition for thermal resources. I am unable to make claims about *E. cyanura* because, of all the individuals observed in hierarchies at high population sites, only 3 of them were *E. cyanura*. This could be a result of bias in my sampling, with most selected sites of observed high populations occurring in areas of closed canopy and at the base of trees. However, there were 6 occasions where 4 or more individuals of *E. cyanura* were basking within 3 meters of one another on the side of an open canopy trail. When I chased them off and the timer was set, none of them returned within the 10-minute period. This may be attributed to the fact that *E. cyanura* prefers open canopy habitats (Bruna et. al 1996; McElroy 2014) and therefore has much more available sunlight in basking habitats. There is less incentive for *E. cyanura* to return to the site of a disturbance if equal amounts of sunlight and similar substrate are available elsewhere, whereas *E. impar* must compete for the sparse sunlight patches coming through the canopy.

Aggressive interactions were observed only between individuals of the same species, and were usually rare and brief. I found nothing to support my hypothesis that this size-based dominance hierarchy allows interspecific exclusion. However, this hierarchy is very much an intraspecific behavior in high competition sites. In most instances, an observed aggressive interaction was a larger *E. impar* pursuing a smaller *E. impar*. Within the hierarchy, smaller individuals were replaced by vacating the substrate, never with aggressive interactions towards the individual replacing them. This may be attributed to the energetic cost of physical and aggressive interactions (Brattstrom 1974). The individual fleeing the encounter behaves submissively because an aggressive encounter would not be energetically efficient.

While more data would be needed to make a claim about *E. cyanura*’s role in a size-based dominance hierarchy, there was a noted trend amongst sites with both species. There were 7 total sites where both species were observed in one 10-minute period. At 5 of these sites, *E. cyanura* was the largest individual present and at one of the other two, *E. cyanura* was within 0.1 mm of the largest individual. It has been proven that *E. impar* moves faster than *E. cyanura* at all temperatures, making *E. impar* a thermal generalist and *E. cyanura* a thermal specialist. (McElroy 2014). *Emoia cyanura* may have to utilize size to out compete the faster *E. impar* when sharing closed canopy habitat. However, in one instance I did observe an *E. impar* sharing a basking substrate with an *E. cyanura*, so competition over sunning substrate is likely not what allows one to exclude the other.

Ultimately, this dominance hierarchy only observed in *E. impar* suggests that, in densely populated locations, intraspecific competition over thermal resources is much more prevalent than interspecific competition. This lack of interspecific competition contributes to their ability to share a habitat, and a size-based dominance hierarchy is not a factor that allows one species to directly exclude another.

For future research, it could be interesting to investigate the relationship between the *Emoia* skinks and *Ficus elastica* trees. The largest populations I observed were in the root network of these large fig trees, with the individuals emerging from within the tree at first light. Similarly, little is known about the home range of individuals. Experimentally, it would be interesting to differentiate individuals by species and sex and observe behavior with experimental crowding.

## ACKNOWLEDGMENTS

Thank you to professors George Roderick, Stephanie Carlson, Seth Finnegan, and Brent Mishler, as well as the incredible Graduate Student Instructors, Phil Georgakakos, Ilana Stein, and Mo Tatlhego. Thank you to the staff of Gump Station, and to my loving classmates who made this semester unforgettable. Lastly, I would like to thank Lady, Skippy, and Taco Bob, who all proved that dogs are truly man’s best friend.

## APPENDIX A

**TABLE 1A.** Location of study sites, elevation, and total number of individuals observed at each site. Species not noted. Some sites are located very close to one another but were each their own 2×2 or 3×3 meter quadrat with no overlap into other sites.

| Site | Latitude | Longitude | Elevation (m) | Total Skinks Observed at Site |
| --- | --- | --- | --- | --- |
| 1 | -149.8297222 | -17.53611111 | 157.9 | 14 |
| 2 | -149.8402778 | -17.52555556 | 38.1 | 25 |
| 3 | -149.8469444 | -17.52055556 | 14.3 | 10 |
| 4 | -149.8341667 | -17.53166667 | 24.1 | 4 |
| 5 | -149.8400000 | -17.52333333 | 32.6 | 15 |
| 6 | -149.8400005 | -17.5234333 | 32.6 | 6 |
| 7 | -149.8347222 | -17.53305556 | 33.8 | 5 |
| 8 | -149.8341668 | -17.53166657 | 23.2 | 8 |
| 9 | -149.8261111 | -17.53666667 | 210.31 | 4 |
| 10 | -149.8277778 | -17.53638889 | 176.8 | 4 |
| 11 | -149.8358333 | -17.54527778 | 213.4 | 8 |
| 12 | -149.8377778 | -17.54555556 | 228.6 | 12 |
| 13 | -149.8436111 | -17.53694444 | 169.8 | 8 |

**FIG. 5A.**
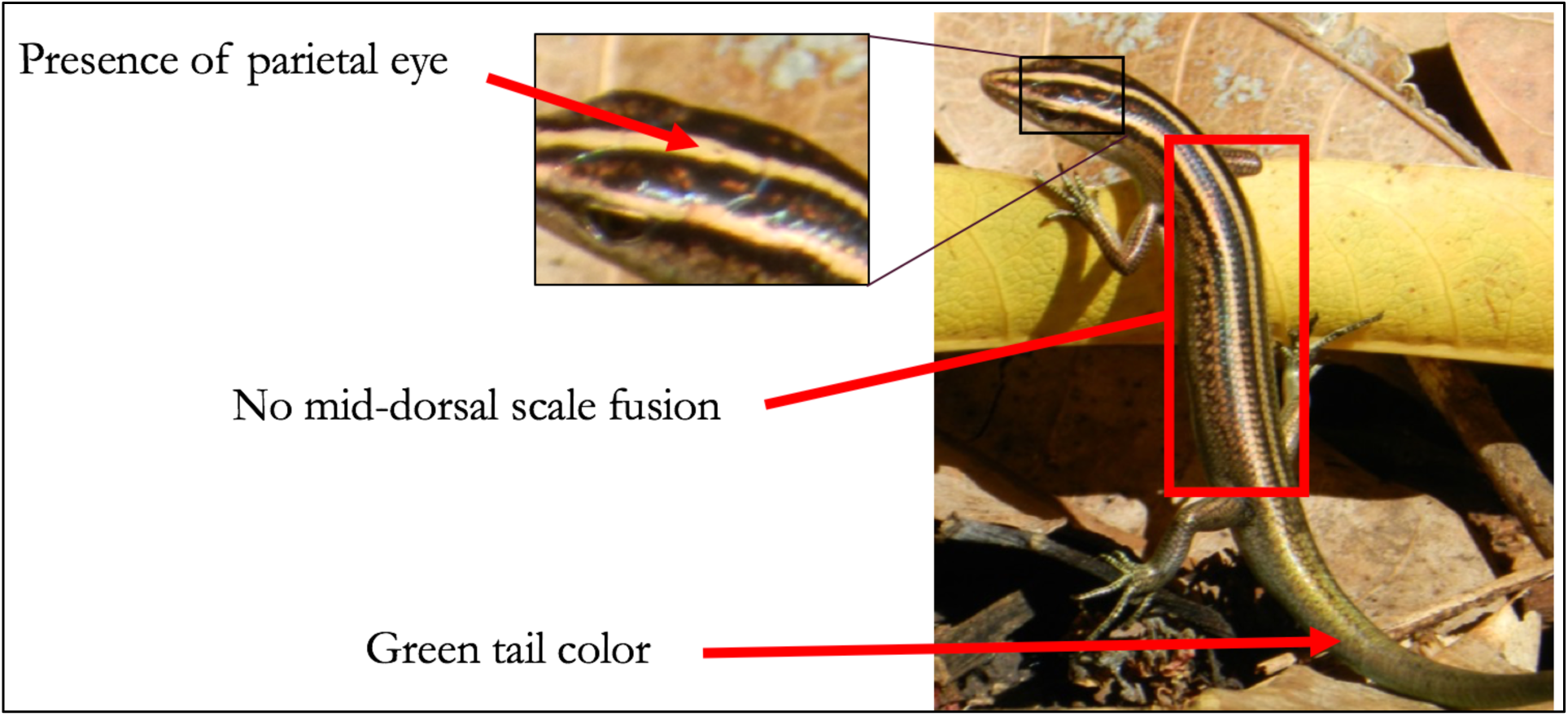
Identification guide for *Emoia cyanura*.

**FIG. 6A.**
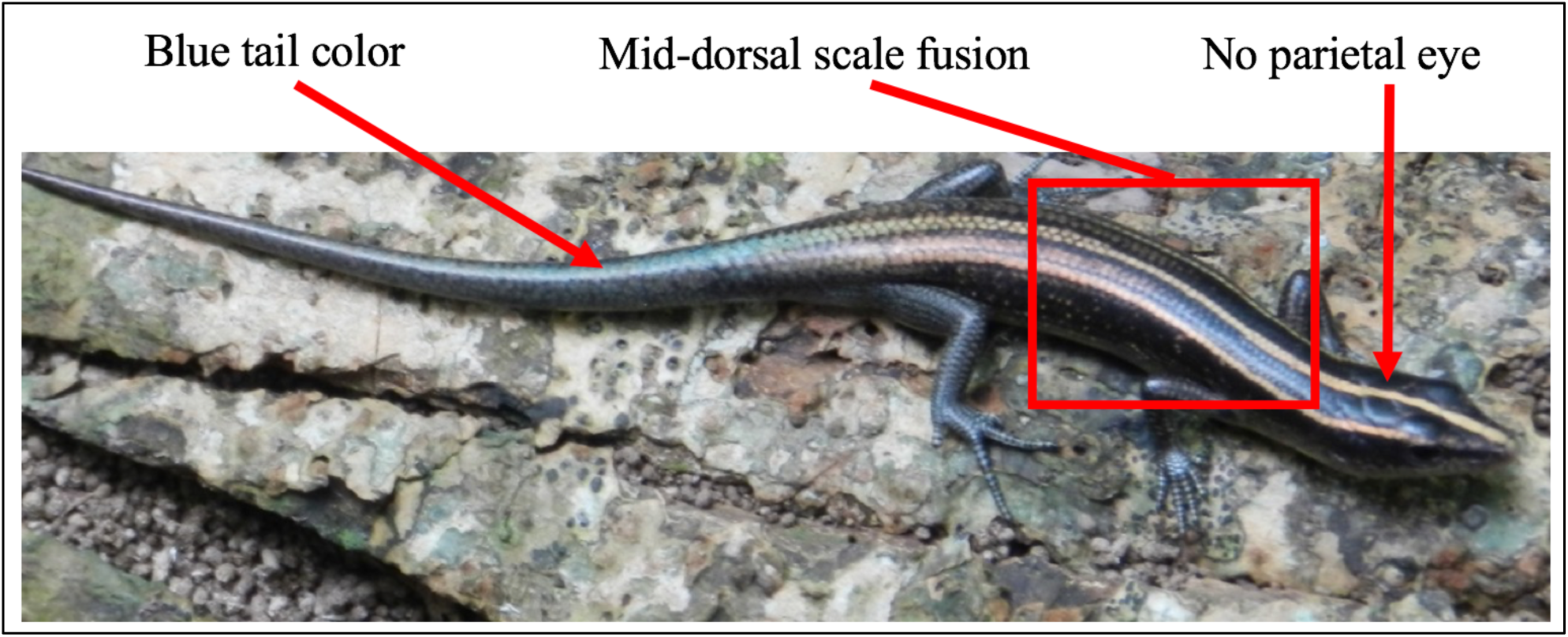
Identification guide for *Emoia impar*.

